# The *Streptococcus agalactiae* R3 surface protein is encoded by *sar5*

**DOI:** 10.1101/2022.01.17.476664

**Authors:** Marte Singsås Dragset, Adelle Basson, Camilla Olaisen, Linn-Karina Selvik, Randi Valsø Lyng, Hilde Lysvand, Christina Gabrielsen Aas, Jan Egil Afset

## Abstract

*Streptococcus agalactiae* (a group B streptococcus; GBS) is an important human pathogen causing pneumonia, sepsis and meningitis in neonates, as well as infections in pregnant women, immunocompromised individuals, and the elderly. For the future control of GBS-inflicted disease, GBS surface exposed proteins are particularly relevant as they may act as antigens for vaccine development and/or as serosubtype markers in epidemiological settings. Even so, the genes encoding some of the surface proteins established as serosubtype markers by antibody-based methods are still unknown. Here, we identify *sar5* as the gene encoding the R3 surface protein, a serosubtype marker of hitherto unknown genetic origin.

## INTRODUCTION

*Streptococcus agalactiae* (a group B streptococcus; GBS) is an important human pathogen, most notably in neonates, but also in pregnant women as well as immunocompromised and elderly individuals. Worldwide, 18 % of pregnant women are colonized with GBS in their rectovaginal tract (1). Colonization of GBS during pregnancy is a risk factor for preterm birth, stillbirth, and neonatal infection (2). To reduce the risk of vertical transmission of GBS to the neonate during birth, routine screening for GBS colonization followed by intrapartum antibiotic prophylaxis (IAP) to pregnant women with GBS is recommended (3). However, administration of IAP poses a risk of allergic and anaphylactic reactions and the widespread use of antibiotics may result in the emergence of antibiotic resistance. Another option to prevent GBS infection is vaccine development. Currently, conserved GBS surface proteins are considered as promising targets for vaccine development (4), as they may elicit a strong immune response against the majority of GBS strains (5).

GBS surface proteins also play an important role as serosubtype markers, relevant for GBS classification in epidemiological settings. While GBS strains can be distinguished into ten serotypes due to differences in their capsular polysaccharide (CPS) (Ia, Ib, and II – IX), surface-expressed protein antigens enable further division of these serotypes. Some of the surface proteins are conserved and present in nearly all GBS strains, while others are associated with specific serotypes, and thus used to define serosubtypes (6). Historically, detection of serosubtypes by means of antibody-based methods has played a major role. In more recent years, serosubtyping of GBS has benefitted greatly from the introduction of molecular methods, such as PCR and whole genome sequencing (WGS) (7, 8).

GBS surface proteins have been classified according to two different and overlapping classification systems (Table 1). However, there is still some discrepancy and confusion surrounding the traditional nomenclature, and some surface proteins that have not yet been definitely linked to a specific gene. One classification scheme of GBS surface proteins includes Cβ and the Cα-like proteins (Alps) Cα, Alp1-4 and Rib. Nearly all GBS strains carry one of the six alp genes (Alp GBS) although, occasionally, an Alp-encoding gene may be absent (non-Alp GBS) (9). Another, and overlapping, classification system of GBS surface proteins is the Streptococcal R proteins first described in 1952 (10), which are resistant to trypsin digestion (thereby designated “R”). R proteins are categorized into five types, R1-5 (11–13). R1 is probably non-existent as a distinct protein; the antiserum raised against R1 was later shown to recognize the identical N-termini of Alp2 and Alp3, the gene products of *alp2* and *alp3*, respectively (14). The R2 protein is expressed by group A and C streptococci and does not seem to occur in GBS (13). The R4 protein has been shown to be identical to Rib and is encoded by the *rib* gene (15), while R5 has been renamed group B protective surface protein (BPS) and was shown to be the gene product of *sar5* (13, 16). The R3 protein has been characterized to some extent (12, 17–19), and has proved useful as a serosubtype GBS marker (20, 21). However, the gene encoding the R3 protein is still unknown (Table 1). BPS was initially thought to be distinct from R3 (13), however, a later study pinpointed a correlation between the presence of the BPS-encoding *sar5* gene and R3 expression (6). Here, we follow up on this correlation, hypothesizing that *sar5* encodes R3. Unraveling the R3-encoding gene, and the putative discrepancy in the nomenclature and nature of the *sar5* gene product, is important for the *sar5* gene product as a prospective target in vaccine development and molecular based GBS serosubtyping, as well as for functional studies on its mechanistic role in pathogenicity.

**Table 1.** Surface-proteins of GBS. Alps (in blue) and R proteins (in green).

| Name | Gene | GenBank Number |
| --- | --- | --- |
| **Cα** | *bca* <sup>(22)</sup> | M97256 |
| **Alp1 (epsilon)** | *alp1* <sup>(23)</sup> | AH013348.2 |
| **Alp2/R1** | *alp2* <sup>(14, 24)</sup> | AF208158 |
| **Alp3/R1** | *alp3* <sup>(14, 24)</sup> | AF245663 |
| **Alp4** | *alp4* <sup>(25)</sup> | AJ488912 |
| **R3** | *unknown* * | - |
| **Rib/R4** | *rib* <sup>(15)</sup> | U583333 |
| **R5/BPS** | *sar5* <sup>(13)</sup> | AJ133114 |

\* in this study determined to be encoded by *sar5*.

## RESULTS

### Presence of the *sar5* gene correlates with R3 protein expression across GBS strains

In a previous study, 121 GBS strains collected from pregnant women in Zimbabwe were tested for (among other markers) the presence of the *sar5* gene and R3 protein expression (6). The study found that 31 out of 35 (91.5%) *sar5* positive strains expressed R3. The remaining 86 strains were negative for both *sar5* and R3. Based on these findings we speculated that *sar5* could encode R3, and that, consequently, the previously reported *sar5*-encoded BPS and the R3 protein are the same protein. To further investigate this observed association between *sar5* and R3 expression, we analyzed 140 clinical GBS strains from neonatal and adult GBS infections from the Norwegian GBS reference laboratory (18). These strains were previously characterized for R3 expression (and other serotype markers) by fluorescent antibody testing using a monoclonal R3 antibody (18). This R3 antibody has been used and evaluated in several previous studies (6, 20, 21, 26–28). We typed the strains for presence/absence of *sar5* using a previously established PCR approach (6), with the R3 reference strain CCUG 29784 (also known as Prague 10/84) as a positive control. Of the 140 GBS strains, the majority were *sar5* negative (131), while nine strains were *sar5* positive. Seven of these strains were R3 positive (Table 2, S1 Figure, and S1 Table). Hence, there was a strong, albeit not perfect, correlation between the presence of the *sar5* gene and R3 expression across the 140 investigated GBS strains.

**Table 2.** Distribution of the *sar5* gene and R3 expression among 140 GBS clinical strains.

|  | **R3 positive** | **R3 negative** | **Sum** |
| --- | --- | --- | --- |
| *sar5* positive | 7 | 2 | 9 |
| *sar5* negative | 0 | 131 | 131 |
| **Sum** | **7** | **133** | **140** |

### The *sar5* positive R3 negative GBS strains express R3 but encode a *sar5* deletion variant

Two strains, 93-33 and 94-3, contained the *sar5* gene but were negative for R3 expression. Thus, these strains were in conflict with our hypothesis that *sar5* encodes R3. The initial detection of R3 in the 140 GBS strains included in this study was performed by fluorescent monoclonal antibody testing on whole bacterial cells (18, 29). We speculated whether R3 from 93-33 and 94-3 could be detected by western blot analysis of denatured proteins from cell lysates. Indeed, using the same R3 monoclonal antibody as in the initial whole cell R3 fluorescence testing, we could clearly see that both 93-33 and 94-3 expressed R3, although in a seemingly truncated form compared to the R3 reference strain CCUG 29784 (Figure 1). For all three strains the blot displayed a ladder-like pattern characteristic for the R3 protein (18). The largest fragment was around the expected size of 109 kDa for the control strain (in accordance with the size of the *sar5* gene at 2940 bp) and around 90 kDa for the 93-33 and 94-3 strains. This finding prompted us to subject GBS strains 93-33 and 94-3 to nanopore WGS, to investigate whether the *sar5* genes of 93-33 and 94-3 could be truncated. Indeed, both GBS strains possessed a *sar5* gene with an identical 531 bp in-frame deletion towards the 3’ end of the gene, when compared to the *sar5* gene of NCTC 9828 (the *sar5* reference strain (13)) (Figure 2). The deletion occurred between two 102 bp long direct repeat regions, making it feasible that the 531 bp region has been deletion by homologous recombination. The 531 bp deletion corresponds perfectly to the 20 kDa difference in size between the R3 control strain and the 93-33 and 94-3 strains observed by western blot analysis (Figure 1). Taken together, our results show that the two *sar5* positive but initially R3 negative strains indeed express R3, although in a truncated form compared to the control strain, and that they both possess a deletion variant of the *sar5* gene.

**Figure 1:**
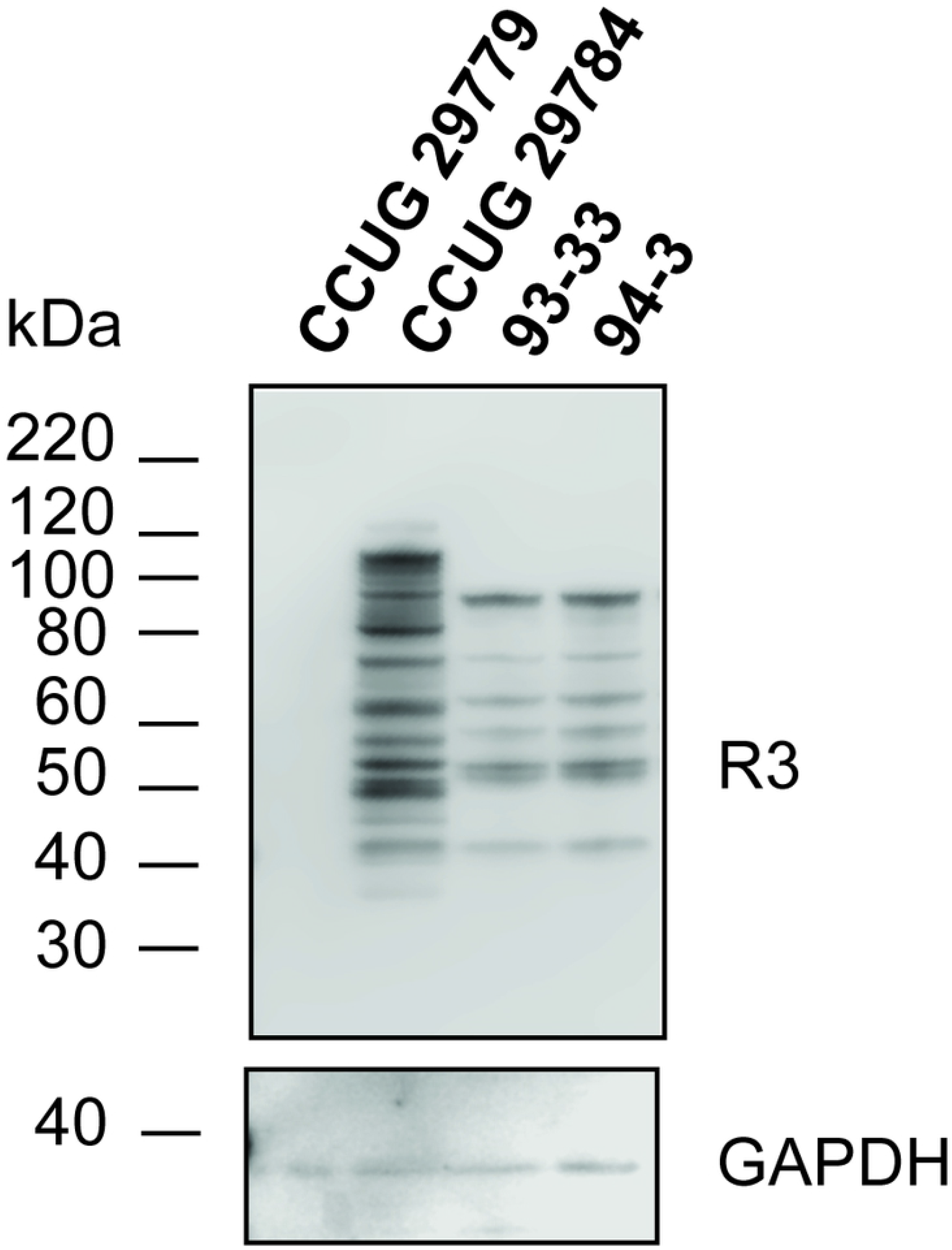
Western blot analysis of GBS whole cell denatured lysates from strains 93-33 and 94-3. The blots were probed with α-R3 antibody (upper panel) and α-GAPDH antibody (lower panel). The CCUG 29779 strain, known to not bind to the R3 antibody, serves as a negative control. The CCUG 29784 strain, known to bind to the R3 antibody, serves as a positive control. The protein standards are shown to the left of the blots.

**Figure 2.**
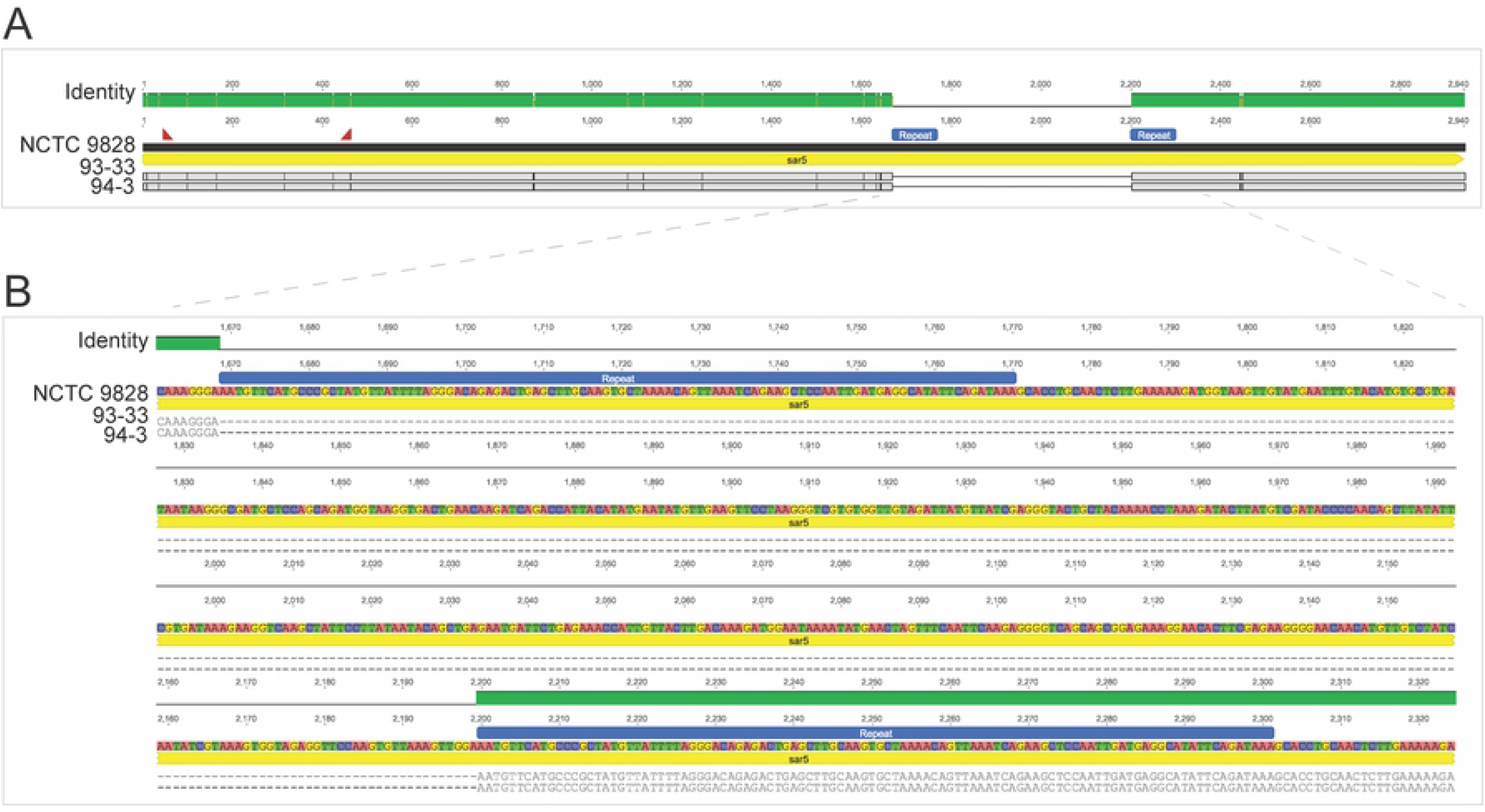
(A) Alignment of the gene sequences of strains 93-33 and 94-3 to the sar5 gene of sar5 reference strain NCTC 9828. Identity between all sequences is indicated by the top panel in green and gene annotation is shown in yellow. Black vertical lines indicate mismatches at the nucleotide level, while grey boxes indicate matching nucleotides to the reference strain. 93-33 and 94-3 have a 531 bp long deletion (marked with horizontal line) within the sar5 gene. The binding sites of the primers used to detect the sar5 gene are shown as red triangles. Repeat regions are shown in blue. (B) Close up of the 531 bp long sar5 deletion in 93-33 and 94-3. The figure is annotated as described for (A) with the additional visualization of sar5 coding strand’s bases.

### The *sar5*-encoded protein is recognized by the R3 antibody

Based on the above results, we had strong indications that the *sar5* gene encodes the R3 protein. We aimed to prove this experimentally by inducing *sar5* protein expression in a *sar5* negative bacterial species, followed by R3 protein detection. First, we constructed a *sar5* inducible expression vector by replacing the luciferase reporter gene of pKT1 (30) with *sar5*, creating pKT1-*sar5*-F. In pKT1, luciferase expression is controlled by the XylS/*Pm* regulator/promoter system, which is induced by the benozoic acid *m*-toluate. We added a FLAG tag to the C-terminal end of the *sar5*-encoded protein, to allow for successful detection of the *sar5-encoded* protein also if the protein was not recognized by the R3 specific antibody (Figure 3). Even so, upon *m*-toluate induction of pKT1-*sar5*-F in *Escherichia coli* BL21 (DE3), we could clearly detect the *sar5*-encoded 109 kDa protein with the monoclonal R3 antibody (Figure 4). We could furthermore detect the FLAG-tag expressed from pKT1-*sar5*-F around the same expected size of 109 kDa, confirming that it is indeed the *sar5* gene product that is detected. When we induced pKT1 (expressing luciferase as opposed to *sar5*), we did not detect expression of any protein around 109 kDa, neither with the R3 nor the FLAG-specific antibodies.

**Figure 3:**
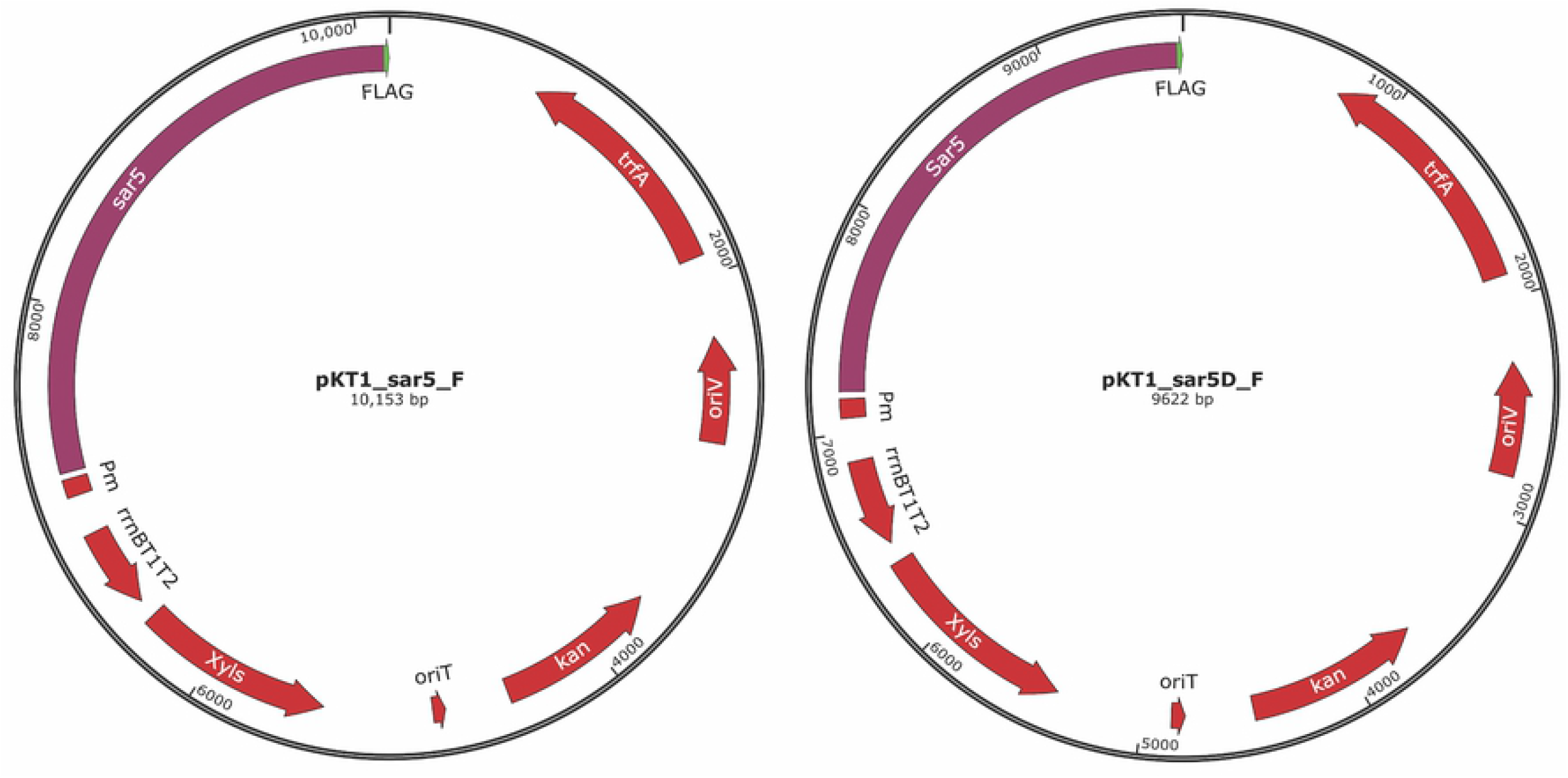
To induce expression of sar5, the luciferase reporter gene of pKT1 (30) was replaced by FLAG-tagged sar5 and FLAG-tagged sar5D, creating pKT1-sar5-F and pKT1-sar5D-F, respectively. xylS, gene encoding the transcription activator XylS. Pm, promoter at which XylS binds and activates transcription in response to the inducer m-toluate. kanR, gene encoding resistance to kanamycin. oriV, origin of replication for RK2-derived plasmids. trfA, gene encoding plasmid replication initiator protein TrfA, activating replication by binding to oriV. rrnBT1T2, transcriptional terminator. oriT, origin of conjugal transfer. The plasmid maps were generated using SnapGene software (from Insightful Science).

**Figure 4:**
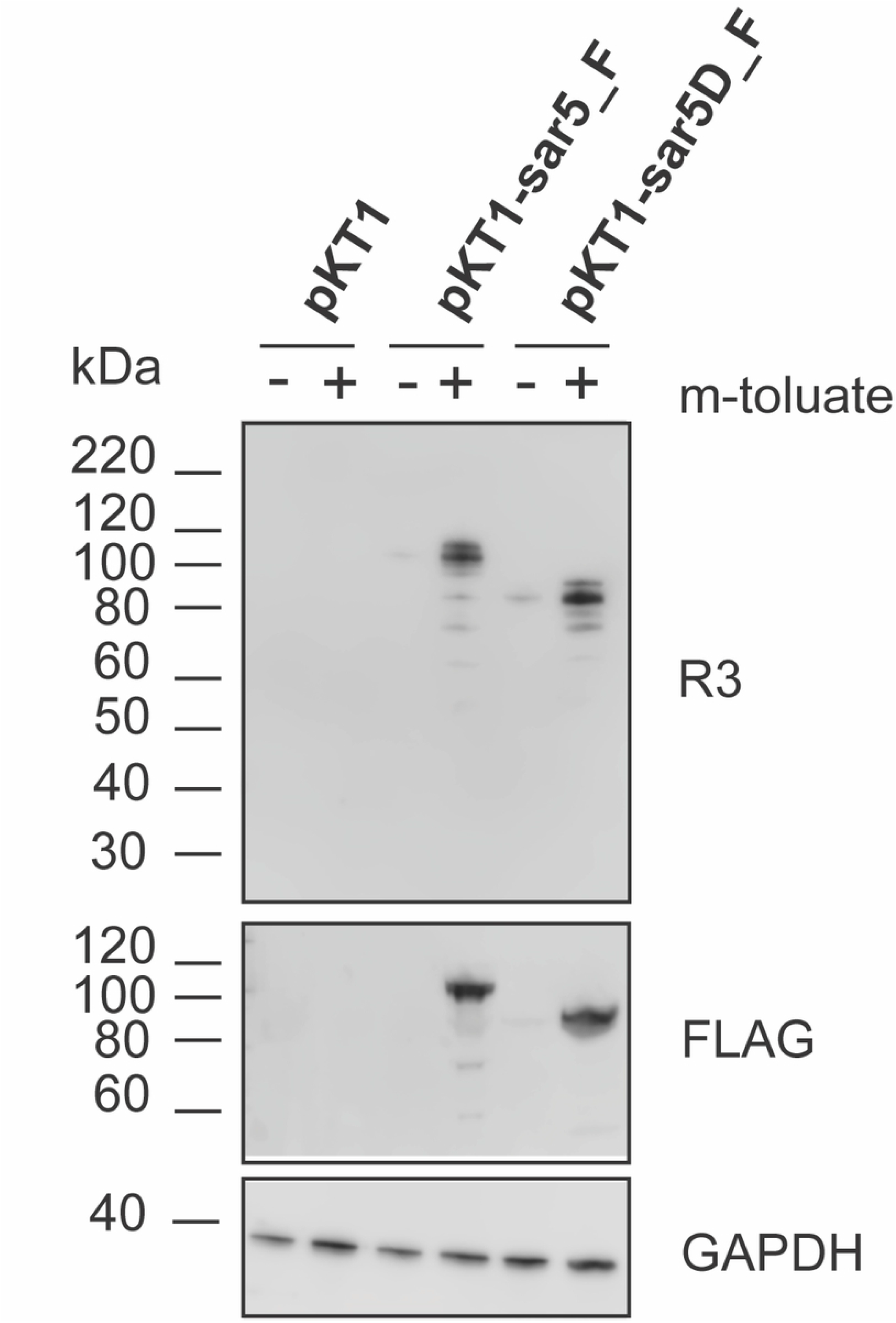
Western blot analysis of whole cell lysates from E. coli BL21 (DE3) carrying pKT1, pKT1-sar5-F or pKT1-sar5D-F plasmids, induced with 2 mM m-toluate (+) or mock-induced with the equivalent amount of the solvent ethanol (-). The blots were probed with r α-R3 antibody (upper panel), α-FLAG antibody (middle panel), or α-GAPDH antibody (lower panel). The protein standards are shown to the left of the blots.

Since the two *sar5* positive but initially R3 negative strains 93-33 and 94-3 actually expressed R3, and were shown to possessed a deletion variant of *sar5* (*sar5D*), we wanted to investigate whether this deletion variant also encoded a protein which is recognized by the R3 antibody. Hence, we constructed an inducible vector expressing FLAG-tagged *sar5D* (pKT1-sar5D-F, Figure 3), and subjected it to induction and western blot analysis. As for the full-length *sar5*, both the R3 and the FLAG antibody bound to the induced *sar5D* gene product (Figure 4). Compared to the full-length *sar5*, the *sar5D* gene encoded a seemingly truncated R3 protein, corresponding in size to the R3 protein expressed by the 93-33 and 94-3 strains. Taken together, our results demonstrate that *sar5* encodes a protein recognized by the R3-specific antibody.

## MATERIALS AND METHODS

### Bacterial strains

Included in this study were 140 clinical GBS strains collected in the years between 1993 and 1995 from neonatal or adult GBS disease from the Norwegian national reference laboratory for GBS, Department of Medical Microbiology, St Olavs Hospital, Trondheim, Norway. These strains were previously characterized for R3 expression (18), as shown in S1 Table. *S. agalactiae* CCUG 29784 was used as an R3 reference strain, while *S. agalactiae* CCUG 29779 (both from Culture Collection University of Gothenburg, Sweden) was used as an R3 negative control in western blot analysis. *S. agalactiae* NCTC 9828 (also called ComptonR, (13)) was used as a reference strain for the *sar5* gene in analysis of *sar5* gene sequences. The GBS strains were cultured overnight on blood agar medium or in Todd-Hewitt broth at 35° C.

### Detection of *sar5* by PCR

Bacterial cells from a single colony were suspended in 100 μl TE-buffer and 100 μl lysis buffer (1% Triton X-100, 0.5% Tween 20, 10 mM Tris-HCl with pH 8 and 1 mM EDTA) (31). The mixture was incubated at 95 °C for 15 minutes and centrifuged at 14 500 rpm for 2 minutes before 100 μl of the supernatant was transferred to a new tube. This material was used as template in the PCR reaction, with AmpliTaq Gold DNA Polymerase with Buffer I (5U/μl; Applied Biosystems). The *sar5*-specific primers used were identical to those of Mavenyengwa et al (6).

### DNA isolation, WGS and assembly

Bacterial cells were suspended in TE buffer and treated with proteinase K (1.5 mg/mL), lysozyme (0.5 mg/mL) and mutanolysin (250 U/mL) for 15 minutes with shaking at 37 °C, before heating at 65 °C for 15 minutes. RNAse A (2 mg/mL) was then added to the lysate. Genomic DNA was subsequently isolated using the EZ1 DNA tissue kit with an EZ1 Advanced XL instrument (Qiagen). Illumina sequencing libraries were prepared using the Nextera XT sample prep kit and sequenced on the Illumina MiSeq platform (Illumina) with 300-bp paired-end read configuration (MiSeq Reagent Kit v3). Nanopore sequencing libraries were prepared using the Rapid Sequencing Kit (SQK-RAD004) and sequenced on a minION intrument with Flongle adapter and flowcells (FLO-FLG001) (Oxford Nanopore Technologies). Raw nanopore data was basecalled using Guppy v5.0.13 and assembled using Flye v2.7. Assemblies were polished with nanopore data using Racon v1.4.20 and with Illumina.data using Pilon v1.23. Geneious vR9 was used for alignments and visualization.

### Cloning of *sar5* into an inducible expression vector

The *sar5* gene was cloned into the *m*-toluate-inducible expression vector pKT1 (30). Briefly, the *sar5* ORF of GBS strain 13/87 (identical to the BPS-encoding gene of strain NCTC 9828 (13)) was amplified. To incorporate a C-terminal FLAG-tag, the sequence encoding the FLAG epitope (DYKDDDDK), was incorporated into the *sar5*-amplification reverse primers. In addition, the primer sets amplifying *sar5* and the pKT1 vector backbone were extended with overlaps to enable Gibson Assembly with the Gibson Assembly^®^ Master Mix (NEB). KOD Xtreme™ Hot Start DNA Polymerase (Sigma-Aldrich) was used for PCR amplifications. Illustrations generated using SnapGene software (Insightful Science; available at snapgene.com) of the cloning strategy and the primers used are found in S1 Materials and Methods.

### Induced expression of *sar5*

For expression of *sar5* in *E. coli*, pKT1 (negative control), pKT1-sar5_F and pKT1-sar5D_F were transformed into *E. coli* strain BL21 (DE3) and grown in LB medium supplemented with 50 μg/ml kanamycin to stationary phase, then diluted approximately 1:500 and grown to OD_600_ 0.05–0.1 at 37 °C. At this point, the samples were adjusted to the same OD_600_, induced with 2 mM of *m*-toluate (Sigma, 1 M stock solution solved in laboratory grade ethanol) and incubated for 5 hours with shaking at 30°C. For uninduced samples the equivalent amount of ethanol was added as a mock treatment.

### Preparation of protein extracts and detection of protein expression by western blot analysis

Overnight cultures of GBS strains were pelleted by centrifugation and washed in PBS. The pellets were resuspended in 1X LDS Sample Buffer (NuPage^®^, Invitrogen) with 50 mM dithiothreitol (DTT) and heated for 10 minutes at 95 °C. Samples were cleared for cellular debris by centrifugation. To prepare protein extracts of *E. coli*, induced (or mock induced) cultures were adjusted to OD_600_ ~0.8, and pelleted by centrifugation. The pellet was resuspended in 50 μl 1x LDS Loading Buffer with 50 mM DTT, heated for 10 minutes at 70°C and sonicated 3 times for 1 minute each. Protein extracts were separated on 4-12% Bis-Tris mini protein gels (NuPage^®^, Invitrogen) and blotted on polyvinylidene fluoride membranes (Bio-Rad). Membranes were blocked with 1X blocking buffer (Roche) in PBS. The primary antibodies against R3 (mouse monoclonal, from (18)), FLAG (monoclonal mouse anti-FLAG M2 antibody, Sigma), and GAPDH antibody GA1R (Covalab) as well as the HRP conjugated secondary antibody goat anti-mouse (Dako) were diluted in 0.5X blocking buffer/PBS. The bound HRP-conjugated antibodies were visualized using SuperSignal™ West Femto Maximum Sensitivity Substrate (Thermo Scientific) and Odyssey Fc imaging system (Licor).

## DISCUSSION

GBS *sar5* was previously shown to encode BPS, a protein initially described to be different from the R3 surface protein (13). Correlation between R3 expression and the presence of the *sar5* gene was observed within a previously examined GBS strain collection, where 31 out of 35 (91.5%) *sar5* positive strains expressed R3 (6). Similarly, frequent co-expression of the Alp protein Cα and the non-Alp Cβ protein has been observed, where 81% of the Cβ positive strains also contained Cα (32). Even so, Cα and Cβ are encoded by two different genes; *bca* and *bac*, respectively (22, 33). To elucidate whether this was also the case for R3 and BPS, we further investigated the correlation between *sar5* and R3 expression across 140 GBS strains from the Norwegian GBS reference laboratory. We observed a perfect correlation between the presence of the *sar5* gene and R3 expression, as well as between the absence of *sar5* and lack of R3 expression. Furthermore, when we induced *sar5* expression in a non-R3 bacterial strain, we found that the monoclonal R3 antibody (18) recognized the *sar5*-encoded protein. During the initial screening of our strain collection two strains were *sar5* positive but R3 negative (93-33 and 94-3). However, while these strains were deemed negative in R3 expression by fluorescent antibody testing on whole bacterial cells (18), they were positive upon western blot analysis of denatured cell lysate (Figure 1). We also found that GBS strains 93-33 and 94-33 possessed a copy of *sar5* with a deletion (*sar5D*). We speculate that the *sar5D-encoded* protein is not recognized by the monoclonal R3 antibody due to conformational changes masking the R3 epitope of the protein in its native form, or that the protein is simply not expressed on the surface of the bacterial cells and thus only detected by immunoblotting of denatured whole cell lysates. Even so, we have demonstrated that the *sar5* gene encodes R3 and that, consequently, R3 and BPS must be one and the same protein.

When BPS was first identified in 2002 by Erdogan *et al*, it was considered a new protein and different from R3 (13). BPS was described in the reference strain NCTC 9828 (called Compton R by Erdogan *et al* (13)), which at that time was considered a Rib and R3 reference strain. However, later that same year, Kong *et al*. reported that the gene thought to encode Rib in strain NCTC 9828 (termed Prague 25/60 by Kong *et al)* had extensive similarities to the *rib* gene but also possessed stretches which differed from *rib* (7). They named this new protein Alp4, which has been the designation used since then (14). Later, it was reported that the C-terminal antigenic determinant of Alp4 and Rib cross-reacted immunologically, while the N-terminal antigenic determinants of Rib and Alp4 differed in immunological specificity (14). The knowledge that NCTC 9828 carries *alp4* (and not *rib*) has consequences for the production of specific antisera targeting R3 and BPS in the study identifying BPS as a distinct protein from R3 (13). Production of specific antibodies was performed by immunizing a rabbit with strain NCTC 9828. Harvested antiserum was adsorbed using a GBS strain expressing *rib*. This would remove antibodies targeting Rib, and is a common procedure for generating specific polyclonal antiserum. However, since NCTC 9828 does not express Rib, but the similar Alp4, antibodies targeting epitopes common to both Rib and Alp4 would be removed by the adsorption whereas antibodies specific only to Alp4 would remain in the antiserum. Immunoprecipitation-bands that were considered evidence of R3 by Erdogan *et al*. in 2002 (13), may in fact have been bands representing Alp4. Similarly, a study reporting 155 (of 4425 total) colonizing and invasive GBS strains expressing BPS found no overlap between R3- and BPS-expression (34). The presumed R3-specific antibody used in that study was also prepared by adsorbing antisera made by immunizing a rabbit with GBS strain NCTC 9828. Regarding BPS and R3 as two distinct proteins would result in an R3-designated antibody without antibodies targeting the *sar5* gene product. Using NCTC 9828 as a reference strain for R3 would thus result in R3 antiserum that may actually detect Alp4.

As a consequence of our findings, already reported data on BPS and R3 are equally relevant for the *sar5* gene product. Using BPS as the future designation of the *sar5* gene product makes the historical R3 protein nonexistent, and vice versa. The nomenclature of GBS surface proteins is already confusing (Table 1). BPS, R5, and now R3 are all names for the same protein. It is important that communication and reports use unambiguous terminology for genes and gene products. We therefore suggest using the designation R3/BPS for the *sar5*-encoded protein henceforth.

Existing data suggest that the prevalence of *sar5* in GBS strains differs between geographical regions. In Norway and the United States, the prevalence has been reported to lie in the range 2.3-8.1 % in invasive GBS strains (18, 27, 34). In a study from the United States, 3.6 % of more than 4000 colonizing GBS strains carried R3/BPS (34), while in Zimbabwe it amounted to near 30 % in healthy pregnant carriers (6). Thus, as a strain variable marker, the R3/BPS protein has proved its potential in serotyping, as a serosubtype marker. Moreover, a recombinant version of this protein has been reported as immunogenic and, on immunization, induced formation of antibodies protective in an animal model, suggesting potential for this protein as a vaccine component (13). R3/BPS may thus be suitable as one of the constituents in a vaccine targeting GBS, particularly in vaccines aimed at populations in areas of Southern Africa (6).

## Acknowledgements

We thank the late Johan A. Mæland for the idea and encouragement for this project.

## Supporting information captions

**S1 Table**: Overview of the R3 typing of the 140 GBS strains examined in this study, as determined by Kvam *et al*. (18), and the *sar5* typing performed in the current study (corresponding to the results of S1 Figure).

**S1 Materials and Methods**: Primers used and illustration of the cloning strategies of pKT1-sar5_F and pKT1-sar5D_F.

**S1 Figure**: The *sar5* typing (PCR amplification) of the 140 GBS strains examined in this study. Negative (-) controls were strain 94-51 (a known R3 negative strain) or H_2_O, positive (+) controls were the R3 reference strain CCUG 29784. A) amplification of the *sar5* gene from the 7 R3 positive GBS strains. B-E) amplification of *sar5* from the 133 R3 negative strains, in pools of 3-5 strains. F) amplification of *sar5* from the strains within pool 7 and pool 9.

